# Selective disruption of synaptic BMP signaling by a Smad mutation adjacent to the highly conserved H2 helix

**DOI:** 10.1101/811109

**Authors:** Tho Huu Nguyen, Tae Hee Han, Stuart Newfeld, Mihaela Serpe

## Abstract

Bone morphogenetic proteins (BMPs) shape normal development and function via canonical and non-canonical signaling pathways. When activating the canonical pathway, BMPs initiate signaling by binding to transmembrane receptors that phosphorylate pathway effectors, the Smad proteins, inducing their translocation into the nucleus and thus regulation of target genes. Phosphorylated Smads also accumulate at cellular junctions, but this non-canonical signaling modality remains less defined. We have recently reported that phosphorylated Smad (pMad in *Drosophila*) accumulates at synaptic junctions in complexes with genetically distinct composition and regulation. Here we examined a wide collection of *Drosophila Mad* alleles and searched for molecular features relevant to pMad accumulation at synaptic junctions. We found that strong *Mad* alleles generally disrupt both synaptic and nuclear pMad accumulation, whereas moderate *Mad* alleles have a wider range of phenotypes and could selectively impact different BMP signaling modalities. Interestingly, synaptic pMad appeared more sensitive to net reduction in Mad levels than nuclear pMad. Importantly, a previously uncharacterized allele, *Mad*^*8*^, showed markedly reduced synaptic pMad levels but only moderately diminished nuclear pMad signals. The postsynaptic composition and electrophysiological properties of *Mad*^*8*^ NMJs were similarly altered. Using biochemical approaches, we examined how single point mutations such as S359L, present in *Mad*^*8*^, could influence the Mad-receptor interface and we identified a key molecular determinant, the H2 helix. Our study highlights the biological relevance of the Smad-dependent, non-canonical BMP signaling and uncovers a highly conserved structural feature of Smads, critical for normal development and function.

## INTRODUCTION

Bone morphogenetic proteins (BMPs) modulate a wide variety of cellular processes via canonical and non-canonical signaling pathways (Derynck and Zhang, 2003; Hogan, 1996; Massague, 1990). Misregulation of BMP signaling is associated with many developmental abnormalities and disease states highlighting the need for tight regulation of various BMP signaling modalities. Like all members of the TGF-β superfamily of signaling factors, BMPs form biologically active dimers that initiate signaling by binding to a heterotetrameric complex of Ser/Thr kinases, known as Type I and Type II BMP receptors (BMPRs). Type II receptors are constitutive kinases that phosphorylate Type I receptors within a regulatory glycine-serine rich (GS) domain to activate them. Activated Type I receptors bind to and phosphorylate the intracellular transducers of the BMP pathway, the R-Smads (Smad1, −5 or −8 in vertebrates and Mad in *Drosophila*) (Feng and Derynck, 2005; Schmierer and Hill, 2007). Phosphorylated R-Smads (pSmads) have a propensity to trimerize that favors their dissociation from the receptors (Kawabata et al., 1998); in the canonical pathway, pSmads associate with co-Smads, translocate into the nucleus and, in conjunction with other transcription factors, modulate expression of target genes.

Activated BMPRs can also signal independently of Smads through noncanonical pathways that include mitogen-activated protein kinase (MAPK), LIM (Lin-11/Isl-1/Mec-3 gene products) kinase, phosphatidylinositol 3-kinase/Akt (PI3K/Akt), and Rho-like small GTPases (Derynck and Zhang, 2003; Moustakas and Heldin, 2005; Zhang, 2009). More recently, pSmad accumulation at cell membranes has been reported in at least two instances: (a) at tight junctions during neural tube closure (Eom et al., 2011), and (b) at the *Drosophila* neuromuscular junction (Dudu et al., 2006; Smith et al., 2012). During neural tube closure, pSmad1/5/8 binds to apical polarity complexes and mediates stabilization of BMP/BMPR complexes at tight junctions (Eom et al., 2011); prolonged BMP blockade disrupts the tight junctions and affects epithelial organization (Eom et al., 2012). In both cases, these Smad-dependent functions do not require transcriptional regulation. In epithelial cells nuclear pSmad signals tend to be very strong, obscuring junctional pSmad signals, making them difficult to study. In contrast, at the fly NMJ, junctional pMad localizes at synaptic terminals, at the contact between motor neurons and body-wall muscles, whereas nuclear pMad accumulates in motor neuron soma and/or nucleus. This spatial separation was instrumental in the initial characterization of this Smad-dependent noncanonical BMP signaling modality (Sulkowski et al., 2014; Sulkowski et al., 2016).

Flies rely on BMP signaling for NMJ growth and neurotransmitter release (Marques and Zhang, 2006). BMP signaling fulfills many of these functions via canonical and noncanonical pathways triggered by Gbb, a BMP7 homolog, which binds to presynaptic BMPRII, Wishful thinking (Wit), and the BMPRIs, Thickveins (Tkv) and Saxophone (Sax). During canonical signaling, these high-order complexes are endocytosed and retrogradely transported to the motor neuron soma where Mad is phosphorylated and regulates transcriptional programs with distinct roles in the structural and functional development of the NMJ. Gbb and Wit also signal non-canonically through the effector protein LIM kinase 1 (LIMK1) to regulate synapse stability (Eaton and Davis, 2005). Importantly, Smad-dependent noncanonical BMP signaling, the third BMP signaling modality, does not require Gbb, but requires Wit, Tkv, Sax, and postsynaptic type-A glutamate receptors (DiAntonio, 2006; Sulkowski et al., 2014). We have previously demonstrated that synaptic pMad is involved in a positive feedback loop across the synaptic cleft: Active, postsynaptic type-A (GluRIIA-containing) glutamate receptors trigger presynaptic accumulation of pMad, which in turn functions to stabilize type-A receptors at postsynaptic densities (Sulkowski et al., 2014; Sulkowski et al., 2016). Genetic and cell biology studies suggest that synaptic Mad is phosphorylated locally by activated BMPRs confined to the active zone, a region with massive exocytosis (of neurotransmitter-filled synaptic vesicles) but no endocytosis. This region effectively traps BMPRs precluding them from endocytosis and participation in canonical BMP signaling. But how does pMad, a product of an enzymatic reaction, remain associated with its own kinase?

To learn more about Mad features that may influence its association with the BMPRs, we have collected most of existing *Drosophila Mad* alleles and compared them side-by side for their ability to sustain the two Smad-dependent signaling modalities: the canonical BMP signaling, marked by pMad accumulation in motor neuron nuclei, and the Smad-dependent noncanonical signaling, marked by pMad accumulation at synaptic terminals. Mad is a highly conserved protein which contains an N-terminal MH1 (Mad homology 1) DNA binding domain, a C-terminal MH2 protein interaction domain, and a linker that has been implicated in the crosstalk with other signaling pathways (Hoodless et al., 1996). Changes in the structure and the oligomeric state of the MH2 domain contribute to the directionality of the signaling process (Wu et al., 2001): On one hand, MH2 mediates Mad binding to BMPRI and thus phosphorylation of its C-terminal SSXS motif, the site of BMP-dependent phosphorylation (Hoodless et al., 1996; Macias-Silva et al., 1996; Zhang et al., 1996). At the same time, MH2 is critical to formation of Smad trimers that dissociate from receptors (Kawabata et al., 1998). In particular, the L3 loop has been implicated in mutually exclusive interactions with the BMPRI and the phosphorylated C-terminal SSXS motif (Wu et al., 2001). Most of molecular lesions in our *Mad* collection map to the MH2 domain, including in the L3 loop, making this allelic series particularly suitable for studies on Mad-Tkv interactions.

Within this comprehensive collection, we found that strong *Mad* alleles generally disrupt both synaptic and nuclear pMad accumulation, whereas moderate *Mad* alleles have a wider range of phenotypes and selectively impact different BMP signaling modalities. In particular, *Mad*^*8*^ showed drastically reduced synaptic pMad levels but only moderately diminished nuclear pMad signals. The postsynaptic composition and electrophysiological properties of *Mad*^*8*^ NMJs were likewise altered. Using biochemical assays and structural modeling, we examined how point mutations such as S359L, present in *Mad*^*8*^, could influence the Mad-Tkv interface. Our study identified a new molecular determinant for this Mad-Tkv interaction, the highly class conserved H2 helix. Several genetic variants identified in human patients map to H2, underscoring the relevance of this motif for normal development and function.

## MATERIALS AND METHODS

### Fly stocks

*Drosophila* stocks used in this study are as follows: *w*^*1118*^, *Mad*^*1*^, *Mad*^*2*^, *Mad*^*3*^, *Mad*^*4*^, *Mad*^*5*^, *Mad*^*6*^, *Mad*^*7*^, *Mad*^*8*^, *Mad*^*9*^, *Mad*^*10*^, *Mad*^*11*^, *Mad*^*12*^ *(Sekelsky et al., 1995)*, *Mad*^*k237*^ (Dworkin and Gibson, 2006), *Mad*^*KG005817*^ (Bellen et al., 2004), *Mad*^*8-2*^ (Wiersdorff et al., 1996), *Df(2L)C28* (Raftery et al., 1995), *twit*^MI06552^ (Venken et al., 2011). The flies were reared at 25°C on Jazz-Mix food (Fisher Scientific). To control for larval crowding, 8 - 10 females were crossed with 5 - 7 males per vial and passed to fresh vials every 3 days.

### Molecular constructs

Flag-Mad plasmid was previously described (Shimmi and O’Connor, 2003). Flag-Mad-8 was generated by PCR followed by Gibson assembly (NEB) and the S359L substitution was verified by sequencing. The PCR primers utilized:

Mad-8-For: 5’-CCAGTACTCGTTCCTCGCCACTCGGAATTCGCGCCCGGTCACTCG;
Mad-8-Rev: 5’-GCGGTATTTTGCACACGGTTAGCGGATGGAATCCGTGGTGG.

To generate the Tac-Tkv construct, the Tac extracellular and transmembrane sequences (*Hin*dIII-*Eco*RI fragment) (Ren et al., 2003) were joined with the PCR amplified wild-type or activated Tkv cytoplasmic domain in the pIB/V5-His vector (ThermoFisher). The activated Tkv chimera (Tac-TkvA) had a single residue substitution Q199D at the end of the GS box (Wieser et al., 1995).

The Tkv primers utilized:

Tkv-For: 5’-GCGTCCTCCTCCTGAGTGGGCTCTGTTTCACCTACAAGCGACGCGAGAAGC;
Tkv-Rev: 5’-GGCTTACCTTCGAACCGCGGGCCCTCTAGACAATCTTAATGGGCACATCG.

### Immunohistochemistry

Larvae were dissected as described previously in ice-cooled Ca^2+^-free HL-3 solution (Budnik et al., 2006; Stewart et al., 1994). The samples were fixed in either 4% formaldehyde (Polysciences) for 30 minutes or in Bouin’s fixative (Bio-Rad) for 3 minutes and washed in phosphate-buffered saline (PBS) containing 0.5% Triton X-100. Primary antibodies from the Developmental Studies Hybridoma Bank were used at the following dilutions: mouse anti-GluRIIA (MH2B), 1:200; rat anti-Elav (7E8A10), 1:200. Other primary antibodies were as follows: rabbit anti-phosphorylated Mad (pMad), 1:500, (a gift from C. H. Heldin), rabbit anti-GluRIIC, 1:1000 (Ramos et al., 2015), Cy5-conjugated goat anti-HRP, 1:1000 (Jackson ImmunoResearch Laboratories, Inc.). Alexa Fluor 488-, Alexa Fluor 568-, and Alexa Fluor 647-conjugated secondary antibodies (Molecular Probes) were used at 1:400. Larval filets and brains were mounted in ProLong Gold (Invitrogen). Samples of different genotypes were processed simultaneously and imaged under identical confocal settings using laser scanning confocal microscopes (CarlZeiss LSM780).

### Fluorescence intensity measurements

Maximum intensity projections were used for quantitative image analysis using Fiji. To measure pMad at the synaptic terminal, synapse surface area was calculated by creating a mask around the HRP signal that labels the neuronal membrane. For each sample, a threshold was applied manually to the HRP channel to remove irrelevant low intensity pixels, outside the NMJ area. The mean puncta intensity was calculated as the total fluorescence intensity signal of the puncta divided their area. All intensities were normalized to control values within an experimental set. The same method was used to quantify synaptic GluRIIA and GluRIIC levels. For nuclear pMad, motor neurons nuclei were labeled with Elav, then thresholds were applied to the Elav signals as above. Mean puncta intensity was calculated as the total fluorescence intensity signal of the puncta divided by the area of the puncta.

### Cell-based assay

A cell-based assay for BMP signaling was described previously (Shimmi and O’Connor, 2003; Serpe and O’Connor, 2006). In brief, S2 cells were transfected with Flag-Mad or 1 Flag-Mad-8. Three days after transfection, cells were incubated with 1nM Dpp (R&D Systems) for 1 h, and cell extracts were analyzed by western blotting. The pMad levels were revealed with anti- phospho-Smad 1/5 (Ser463/465) 41D10 (Cell signaling) 1:200 and anti-Tubulin (Invitrogen) 1:500, and quantified relative to the Flag signal detected with anti-Flag M2 (Sigma) at 1:2,000 using IRDye secondary antibodies for simultaneous detection on an Odyssey Infrared Imaging System (Li-Cor Biosciences).

For immunocytochemistry experiments, eight-well chambers (Fisher scientific) were coated with 0.1 mg/ml Concanavalin A (Sigma) for 1 hr at 25°C, then with 1 ng/ml anti-Tac antibodies (M-A251, BioLegend) for 1 h at 25°C. Transiently transfected S2 cells were grown for three days before spreading on ConA/anti- Tac- coated chamber for 1 h at 25°C. The surface attached cells were fixed in 4% formaldehyde (Polysciences) for 15 min, then stained for pMad (anti-phospho-Smad 1/5, 41D10, 1:200, Cell Signaling), and Flag (anti-Flag, M2, 1:500, Sigma). FITC- conjugated goat anti-phalloidin, 1:500 (Invitrogen), Alexa Fluor 568-, and Alexa Fluor 647-conjugated secondary antibodies (Molecular Probes) were used at 1:400. Cells were mounted in DAPI-containing Prolong Gold (Invitrogen).

### Electrophysiology

Recordings were performed on muscle 6, segment A3 of third instar larvae as previously reported (Qin et al., 2005). Briefly, wandering third instar larvae were dissected in ice-cold, calcium-free physiological HL-3 saline (Stewart et al., 1994) and immersed in HL-3 containing Ca^2+^ before being shifted to the recording chamber. The calcium-free HL-3 saline contains (in mM): 70 NaCl, 5 KCl, 20 MgCl_2_, 10 HCO_3_, 5 trehalose, 115 sucrose, 5 HEPES, pH adjusted to 7.2 at room temperature. The recording solution was HL-3 with either 0.5 mM CaCl_2_. Intracellular electrodes (borosilicate glass capillaries of 1 mm diameter) were filled with 3 M KCl and resistances ranged from 12 to 25 MΩ. Recordings were done at room temperature from muscle cells with an initial membrane potential between −50 and −70 mV, and input resistances of ≥ 4 MΩ. For mEJC recordings the muscle cells were clamped to −80 mV. To calculate mean amplitudes and frequency of mEJCs, 100–150 events from each muscle were measured and averaged using the Mini Analysis program (Synaptosoft). Minis with a slow rise and falling time arising from neighboring electrically coupled muscle cells were excluded from analysis (Gho, 1994; Zhang et al., 1998). To compare decay time constant of mEJCs between genotypes, 50 clear representative events from each recording were averaged and fitted using a double exponential equation of the form

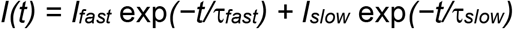

where *I*_*x*_ is the peak current amplitude of a decay component and τ_x_ is the corresponding decay time constant. To allow for easier comparison of decay times between genotypes weighted τ (ms) were calculated using the formula

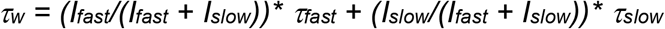

Electrical signals were recorded with an Axoclamp 2B amplifier (Axon Instruments). The signals were filtered at 1 kHz and digitized at 10 kHz by using an analog-digital converter (Digidata 1440A) and pCLAMP software (version 10.0, Axon Instruments). Data are presented as mean ± SEM. Two-tailed unpaired Student’s *t* test was used to assess statistically significant differences among genotypes. Differences were considered significant at *p* < 0.05.

## RESULTS

### Analysis of pMad in larval motor neurons in distinct *Mad* alleles

To test the effect of *Mad* mutations on the distribution and levels of synaptic and nuclear pMad, 15 available *Mad* alleles were examined (Table 1). Most of these alleles have been isolated as dominant maternal enhancers of recessive *dpp* alleles (Chen et al., 1998; Raftery et al., 1995; Sekelsky et al., 1995). We crossed each of these alleles with *Df(2L)C28*, a small deficiency covering the *Mad* locus, counted the resulted progenies, and calculated the percent of expected adult progeny that were trans-heterozygous for the allele and the deletion (Table 1). As expected, we observed only a few *Mad*^*i*^/*Df(2L)C28* adult escapers, with the largest escaper percentage from the weak *Mad*^*5*^ and *Mad*^*6*^ alleles, previously classified as hemizygous viable (Sekelsky et al., 1995). In all crosses we observed larval and pupal lethality as *Mad*-deficient animals were unable to enter and/or complete metamorphosis. The *Mad*^*i*^/*Df(2L)C28* larvae, henceforth referred to as “*Mad*^*i*^ mutants”, have developmental delays and a transparent appearance due to reduced fat body. Since BMP signaling, primarily through Gbb/BMP7, is a central player in the energy homeostasis (Ballard et al., 2010), we used fat body accumulation as an additional metric for the severity of *Mad* mutant phenotypes (below).

**Table 1.**
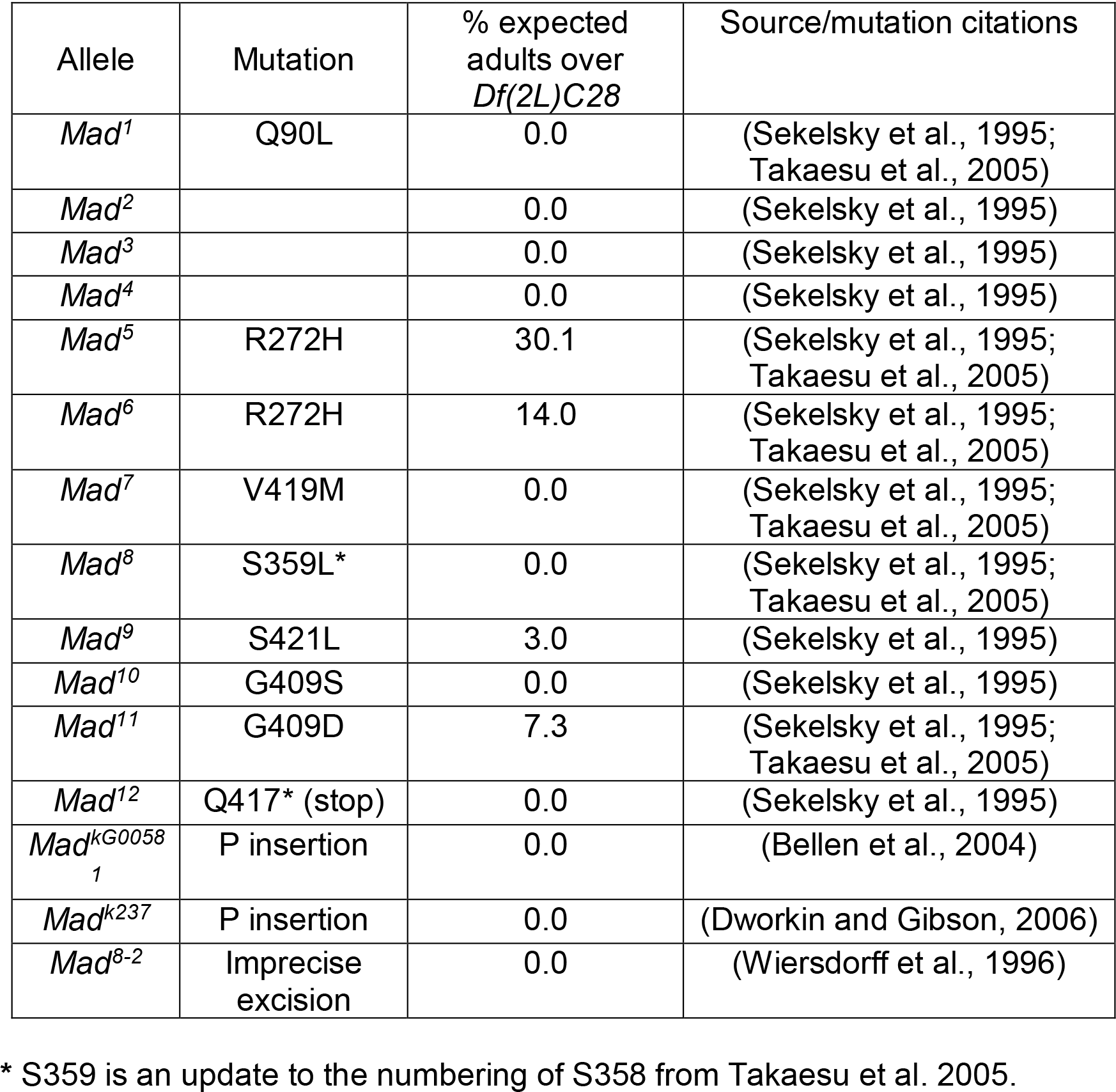
List of *Mad* alleles analyzed with viability data

To quantify the accumulation of pMad in motor neuron nuclei, we co-stained third instar larval ventral ganglia for pMad and Elav, a neuronal nuclear factor. For synaptic pMad, we quantified the pMad NMJ signals relative to anti-HRP, which labels neuronal membranes (Jan and Jan, 1982). We used as negative control the well characterized *Mad*^*12*^ mutant to examine any residual pMad staining at the larval body wall muscle NMJ and in motor neurons (Figure 1A-B). The protein encoded by *Mad*^*12*^ allele has a C-terminal truncation (Q417* stop) prior to the BMP signaling specific phosphorylation site, S^451^SXS, that abolishes any zygotic pMad signals (Sekelsky et al., 1995). In *Mad*^*12*^ mutants, synaptic and nuclear pMad were reduced by 80 ± 5% and 91 ± 5%, respectively (Table 2). The residual signals may be due to (a) maternal wild-type Mad protein, that may be present in very low amounts during third instar larval stages, (b) cross reaction of the pMad antibodies with pSmox, the R-Smad of the Activin signaling pathway in *Drosophila*, that shares high homology with Mad over the C-terminus, or (c) background staining.

**Table 2.** Summary of synaptic and nuclear pMad levels

| Genotype | Synaptic pMad |  |  |  | Nuclear pMad |  |  |  |
| --- | --- | --- | --- | --- | --- | --- | --- | --- |
|  | Mean | SEM | n | P value vs control | Mean | SEM | n | P value vs control |
| Control | 100 | 2.473 | 30 | 1 | 100 | 2.0813 | 29 | 1 |
| *Mad*<sup>12</sup>/*Df* | 19.7084 | 5.25 | 18 | 4.243E-11 | 9.3579 | 5.19 | 17 | < 2.2E-16 |
| *Mad*<sup>7</sup>/*Df* | 23.5568 | 4.439 | 21 | 1.698E-10 | 13.2623 | 4.854 | 14 | < 2.2E-16 |
| *Mad*<sup>10</sup>/*Df* | 22.8839 | 4.511 | 23 | 1.331E-11 | 20.49043 | 3.6048 | 15 | < 2.2E-16 |
| *Mad*<sup>4</sup>/*Df* | 24.0696 | 1.5151 | 17 | < 2.2E-16 | 7.8218 | 2.3397 | 12 | < 2.2E-16 |
| *Mad*<sup>5</sup>/*Df* | 99.0553 | 5.38 | 14 | 0.8344 | 97.5586 | 4.6382 | 17 | 0.6252 |
| *Mad*<sup>6</sup>/*Df* | 88.2728 | 2.1829 | 14 | 0.8701 | 93.67356 | 4.4325 | 13 | 0.7717 |
| *Mad*<sup>k237</sup>/*Df* | 63.9902 | 3.01328 | 41 | 1.585E-11 | 75.3151 | 3.35743 | 19 | 1.31E-06 |
| *Mad*<sup>k237</sup>/*Mad*<sup>k237</sup> | 58.5808 | 3.01725 | 36 | 1.47E-13 | 60.0307 | 4.20185 | 17 | 9.42E-09 |
| *Mad*<sup>8-2</sup>/*Df* | 50.2349 | 4.401 | 21 | 2.42E-11 | 57.35443 | 4.115434 | 16 | 3.1E-09 |
| *Mad*<sup>9</sup>/*Df* | 62.9174 | 4.4278 | 17 | 1.516E-07 | 79.93903 | 2.008 | 16 | 1.92E-05 |
| *Mad*<sup>11</sup>/*Df* | 61.7551 | 1.4798 | 19 | 1.398E-11 | 74.02319 | 4.9773 | 13 | 0.000199 |
| *Mad*<sup>1</sup>/*Df* | 45.981 | 2.8567 | 19 | 5.63E-10 | 31.62352 | 5.0572 | 15 | 1.02E-08 |
| *Mad*<sup>2</sup>/*Df* | N/A |  |  |  | N/A |  |  |  |
| *Mad*<sup>3</sup>/*Df* | N/A |  |  |  | N/A |  |  |  |
| *Mad*<sup>KG00581</sup>/*Df* | N/A |  |  |  | N/A |  |  |  |
| *Mad*<sup>8</sup>/*Df* | 53.5168 | 1.6756 | 22 | 3.44E-09 | 75.42566 | 3.2583 | 15 | 1.42E-12 |

**Figure 1.**
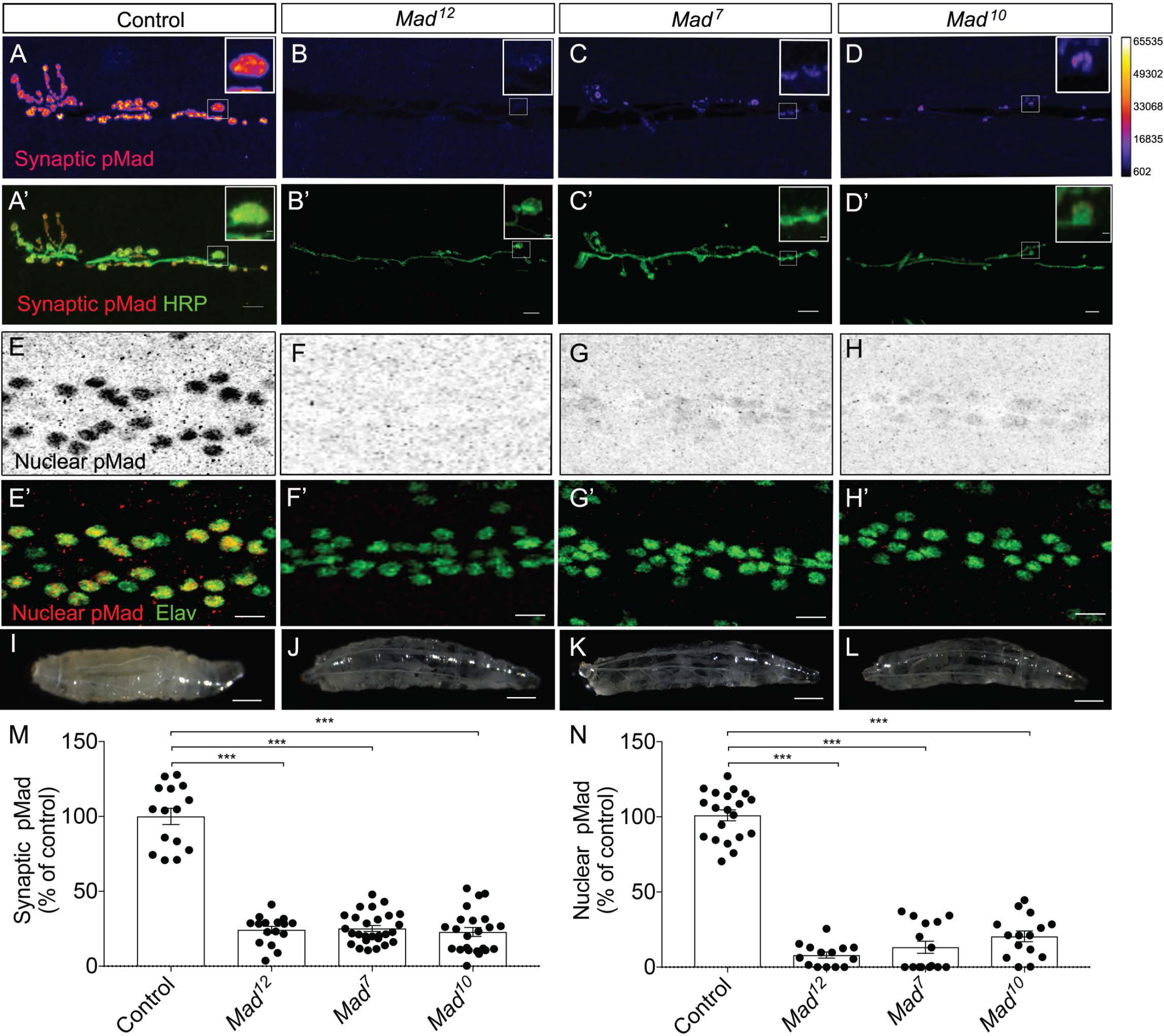
Strong Mad alleles disrupt both synaptic and nuclear pMad. (A-D) Representative confocal images of NMJ 6/7 boutons from third instar larvae of indicated genotypes labeled for pMad (red) and HRP (green). Synaptic pMad is showed in Fire-lut representation in the upper panels; on the intensity scale, white represents peak intensity (~6,000 arbitrary units, A.U.). (E-H) Confocal images of ventral nerve cord of larvae of indicated genotypes labeled with pMad (red in the merged panels) and Elav (green). In comparison with control animals (E), nuclear pMad is strongly reduced in *Mad*^*12*^ (F), *Mad*^*7*^ (G), and *Mad*^*10*^ (H) mutants. (I-L) Third instar control larvae (I) have opaque appearance, while *Mad* mutants (J-L) are almost transparent due to severely reduced fat body. (M-N) Quantitation of synaptic (M) and nuclear pMad (N) levels. See also Table 2. Scale bars: 10 μm (A-H) and 1 μm (details); 100 μm (I-L). Error bars indicate SEM. ***, p <0.0001.

### Strong *Mad* alleles drastically reduced both synaptic and nuclear pMad signals

*Mad*^*7*^ and *Mad*^*10*^ are strong alleles with single residue substitutions within the MH2 domain: V419M in *Mad*^*7*^ and G409S in *Mad*^*10*^, respectively (Sekelsky et al., 1995; Takaesu et al., 2005). Both residues map to the L3 loop of the MH2 domain, which has been implicated in mutually exclusive interaction with the type I receptors or with the phosphorylated pS-X-pS tail of adjacent Smads in trimeric complexes (Wu et al., 2001). As expected, both *Mad*^*7*^ and *Mad*^*10*^ mutants exhibited a drastic reduction of synaptic and nuclear pMad (Figure 1C-D, quantified in M-N). Compared to the *w*^*1118*^ control, the synaptic pMad levels were reduced to 24 ± 4% in *Mad*^*7*^ and to 23 ± 4% in *Mad*^*10*^; such levels were very close to those observed in the negative control, *Mad*^*12*^ (Figure 1B and Table 2). The nuclear pMad signals were similarly reduced to 13 ± 5% and 20 ± 5% in *Mad*^*7*^ and *Mad*^*10*^ respectively. The *Mad*^*7*^, *Mad*^*10*^ and *Mad*^*12*^ mutant larvae appear translucent, with strongly reduced fat body, indicating severely impaired canonical BMP signaling (Figure 1 I-L).

*Mad*^*7*^, *Mad*^*10*^ and *Mad*^*12*^ were previously classified as null alleles based on their maternal effect enhancement of *dpp* (Sekelsky et al., 1995). Our analyses indicate that these alleles also behave as very strong hypomorphs during larval stages. In contrast, *Mad*^*4*^ was initially classified as moderate allele; however, we found that both synaptic and nuclear pMad levels were drastically reduced in the *Mad*^*4*^ mutant larvae to 24 ± 2% and 8 ± 2% respectively, compare to control (Table 2). This result could not be explained by additional lesions on *Mad*^*4*^ chromosome, as we analyzed *Mad*^*4*^/*Df(2L)C28* trans-heterozygote animals. Rather, we speculate that the unidentified molecular lesion in the *Mad*^*4*^ allele leaves out critical function(s) during larval development.

At the opposing pole, the weak *Mad*^*5*^ and *Mad*^*6*^ mutants showed no significant changes in either synaptic or nuclear pMad levels during larval stages (Table 2) and had normal larval fat body (not shown). These alleles were isolated independently but have the same single residue change, R272H (Sekelsky et al., 1995; Takaesu et al., 2005). Nonetheless, these mutants yielded few adult escapers (Table 1), indicating that the substitution of this residue triggers critical deficits during later developmental stages.

### Moderate *Mad* alleles exhibited differential effects on synaptic and nuclear pMad

While single point mutations disrupt specific functional domains within Mad protein, the *Mad*^*K237*^ allele (or *Mad^k00237^*) has reduced overall Mad expression due to the insertion of a transposomal element within the *5’UTR* of *Mad mRNA* (Dworkin and Gibson, 2006). Previous studies reported significant NMJ defects in *Mad*^*K237*^ homozygous and trans-heterozygote combinations (Merino et al., 2009). We also found disruptions of pMad levels in *Mad*^*K237*^ third instar larvae (Figure 2A-B and Table 2). Interestingly, the pMad levels were differentially disrupted in *Mad*^*K237*^ mutants, with the synaptic pMad reduced to 64 ± 3% and the nuclear pMad to 75 ± 3% compare to control. A separate *Mad* allele, *Mad*^*8-2*^, with a lesion in the first intron of the *Mad* gene, showed very similar behavior (Table 2). This suggests that synaptic pMad is more sensitive to suboptimal Mad levels than the nuclear pMad. A similar trend was observed for other moderate *Mad* alleles, such as *Mad*^*9*^ and *Mad*^*11*^, which code for single residue substitutions, S421L and G409D, respectively, in the L3 loop of the MH2 domain (Sekelsky et al., 1995; Takaesu et al., 2005). In these mutants, the synaptic pMad was reduced to 63 ± 4% and 62 ± 4%, respectively, while nuclear pMad was reduced to only 80 ± 4% and 74 ± 4% compared to control (Figure 2C-D, quantified in M-N). A relatively small reduction of canonical BMP signaling was also evident when examining the larval fat body: These *Mad* mutants had modestly reduced fat body in comparison with the control (Figure 2I-L). The synaptic pMad deficits observed in moderate *Mad* alleles suggest that non-canonical BMP signaling is sensitive to Mad net levels and to the integrity of the Mad-Type-I receptor interface, including the L3 loop.

**Figure 2.**
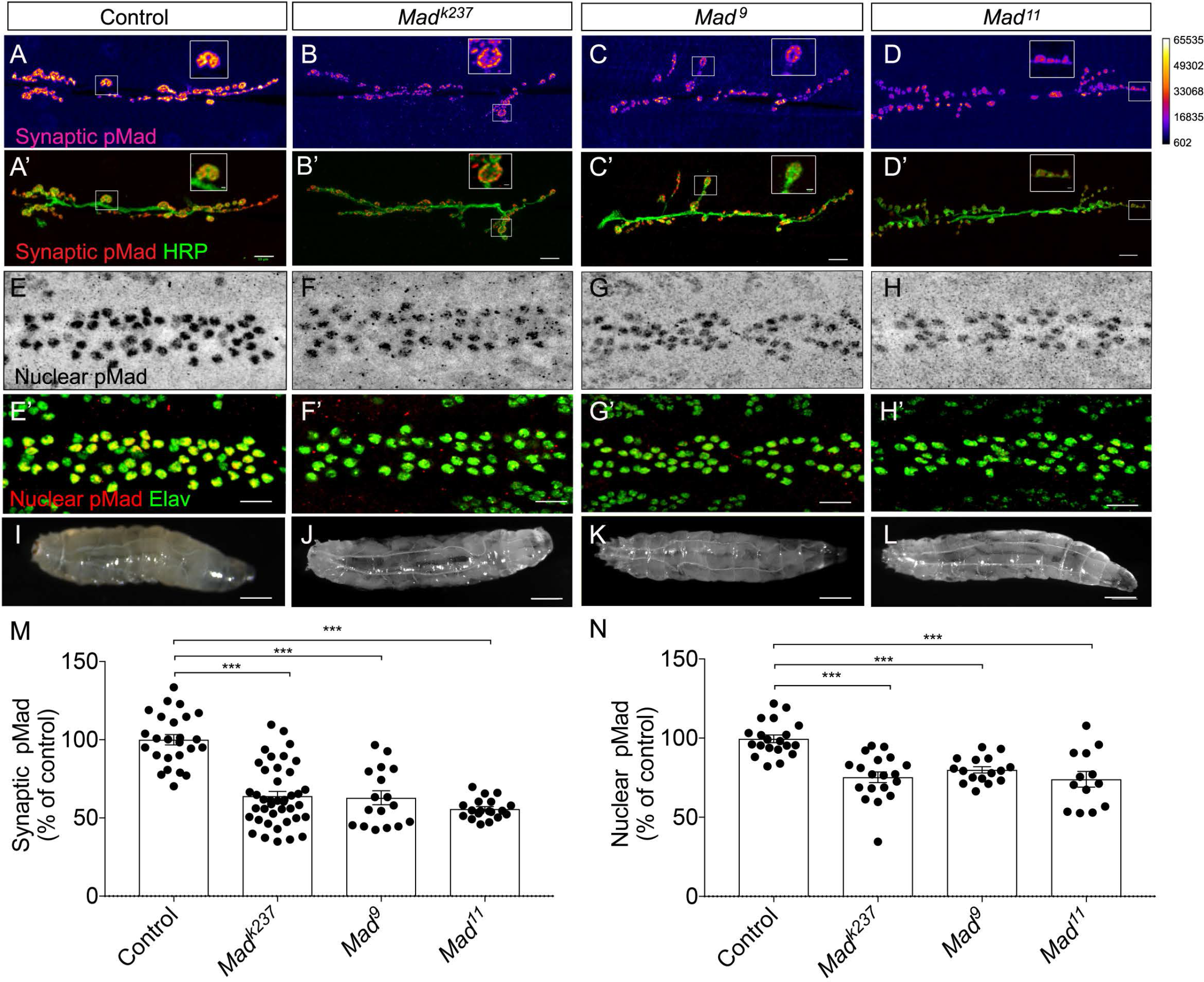
Moderate *Mad* alleles exhibit differential effects on synaptic and nuclear pMad. (A-H) Confocal images of NMJ 6/7 boutons and ventral nerve cord from third instar larvae of indicated genotypes labeled for pMad (red) and HRP (green) (A-D) or Elav (green) (E-H). Synaptic pMad is showed in Fire-lut in the upper panels; on the intensity scale, white represents peak intensity (~6,000 arbitrary units, A.U.). (E-H) Confocal images of ventral nerve cord of larvae of indicated genotypes labeled with pMad (red in the merged panels) and Elav (green). (I-L) Images of third instar larvae. *Mad* mutants (J-L) appear slightly translucent in comparison with control (I) due to mildly reduced fat body. (M-N) Quantitation of synaptic (M) and nuclear pMad (N) levels. See also Table 2. Scale bars: 10 μm (A-H) and 1 μm (details); 100 μm (I-L). Error bars indicate SEM. ***, p <0.0001.

### *Mad*^*1*^ – the only allele with a predominant effect on nuclear pMad

*Mad*^*1*^ is the only known *Drosophila Mad* allele with a mutation in the DNA binding domain of Mad, (Q90L within the MH1) (Takaesu et al., 2005). *Mad*^*1*^ has been classified as a gain-of-function allele with dominant-negative activity because, in comparison with a *Mad* deletion, it exhibited an enhanced effect on *dpp*^*s6*^/*dpph*^*r4*^ vein phenotypes (Takaesu et al., 2005). Interestingly, *Mad*^*1*^ was one the few alleles in our study with relative reduction of nuclear pMad signals exceeding that of synaptic pMad: in *Mad*^*1*^ mutants, the pMad levels were reduced to 46 ± 3% and 32 ± 5% of control synaptic and nuclear pMad, respectively (Figure 3A-D). This likely reflects the molecular nature of this mutation, which may disrupt Mad binding to DNA more than Mad interacting with other proteins. The fat body was also strongly reduced in the *Mad*^*1*^ mutant, consistent with *Mad*^*1*^ severe canonical BMP signaling defects (Figure 3E-F).

**Figure 3.**
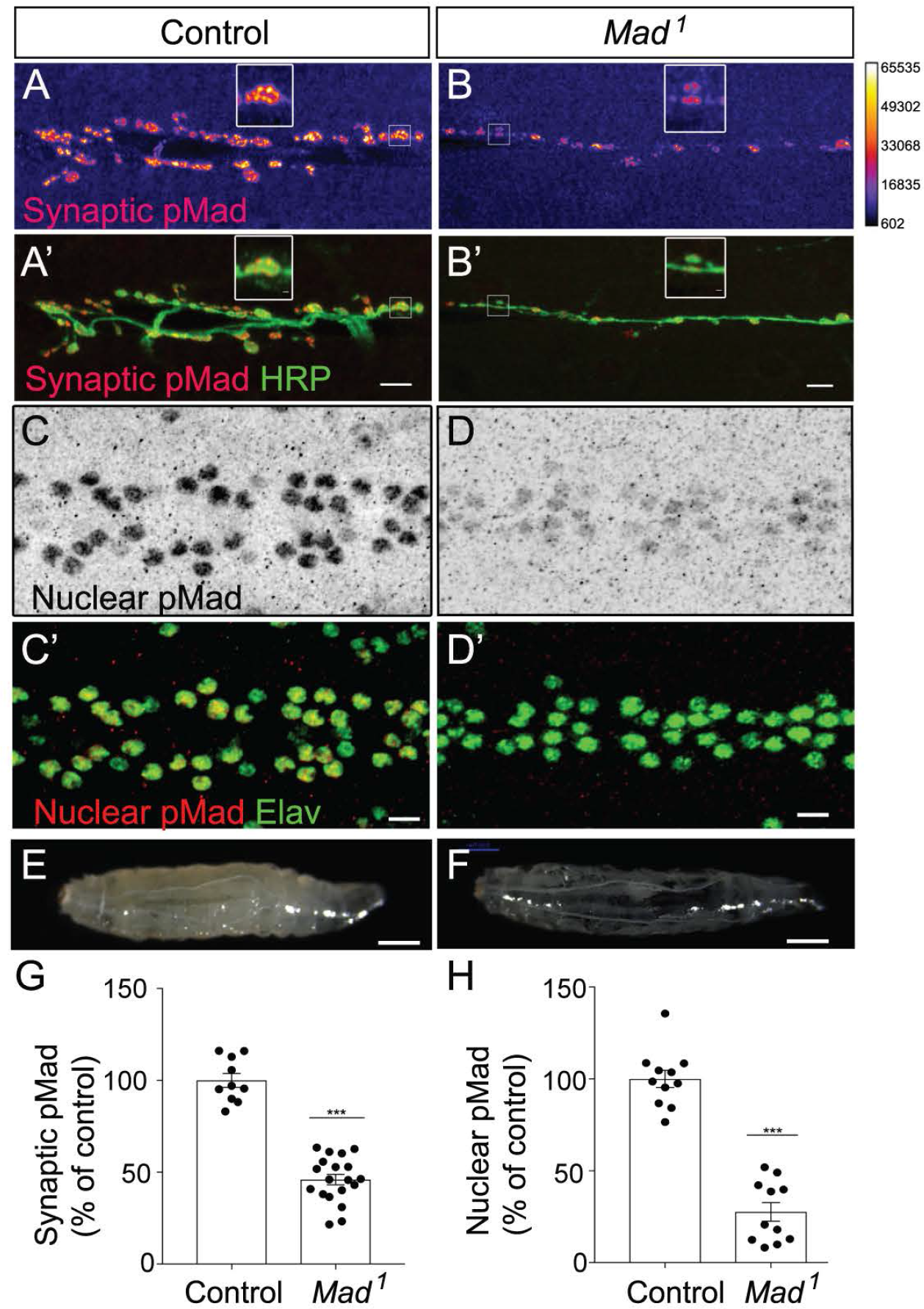
A single residue change in Mad DNA binding domain induces predominantly nuclear pMad deficits. (A-D) Confocal images of third instar NMJ 6/7 boutons and ventral nerve cord from control and *Mad*^***1***^ mutants labeled for pMad (red) and HRP (green) (A-B) or Elav (green) (C-D). (E-F) The level of fat body is strongly reduced in *Mad*^*1*^ mutants (F) in comparison to control (E). pMad is showed in Fire-lut in the upper panels (A-B) and is quantified in (G-H). Scale bars: 10 μm (A-D) and 1 μm (details); 100 μm (E-F). Error bars indicate SEM. ***, p <0.0001.

In our analyses, *Mad*^*2*^ and *Mad*^*3*^ alleles appeared more detrimental than *Mad*^*1*^, *Mad*^*7*^, *Mad*^*10*^ and *Mad*^*12*^: *Mad*^*2*^ (or *Mad*^*3*^)/*Df(2L)C28* animals died before reaching the third instar stage, whereas all other *Mad* mutants we examined exhibited only partial lethality as third instar larvae. A similar result was obtained for *Mad*^*KG00581*^, a strong allele generated by the insertion of a transposon element within the *Mad* gene (Bellen et al., 2004). We did not characterize these strong *Mad* alleles further. Instead, in search for molecular determinants relevant for pMad accumulation at synaptic junctions, we turned our attention towards alleles that differentially modulate different BMP signaling modalities.

### *Mad*^*8*^ exhibited a disproportionate reduction of synaptic pMad

Aside from *Mad*^*K237*^ and the loop L3 mutants, most *Mad* alleles showed a relatively proportional reduction in both synaptic and nuclear pMad levels (Table 2). A notable exception was *Mad*^*8*^, which codes for protein with a single residue change, S359L, outside of any functional motif of Mad (Takaesu et al., 2005). *Mad*^*8*^ exhibited a very significant reduction in synaptic pMad (to 54 ± 5% compared to control) but a more modest reduction in nuclear pMad (to 75 ± 5%) (Figure 4A-D, Table 2). Moreover, there was no readily apparent difference in the extent of the fat body in *Mad*^*8*^ mutants in comparison to control third instar larvae (Figure 4E-F). We further confirmed the small effect of the *Mad*^*8*^ allele on nuclear pMad signals by examining the motor neurons expression of *twit (<u>t</u>arget of <u>wit</u>)*, a gene regulated by BMP signaling (Kim and Marques, 2010). Using a *twit∷GFP* insertion (Sulkowski et al., 2016; Venken et al., 2011), we found that Twit∷GFP levels were reduced to 70 ± 4% in the *Mad*^*8*^ mutant motor neurons in comparison to control (Figure 4I-K). These data are consistent with a relatively moderate reduction of nuclear pMad in *Mad*^*8*^ larval motor neurons.

**Figure 4.**
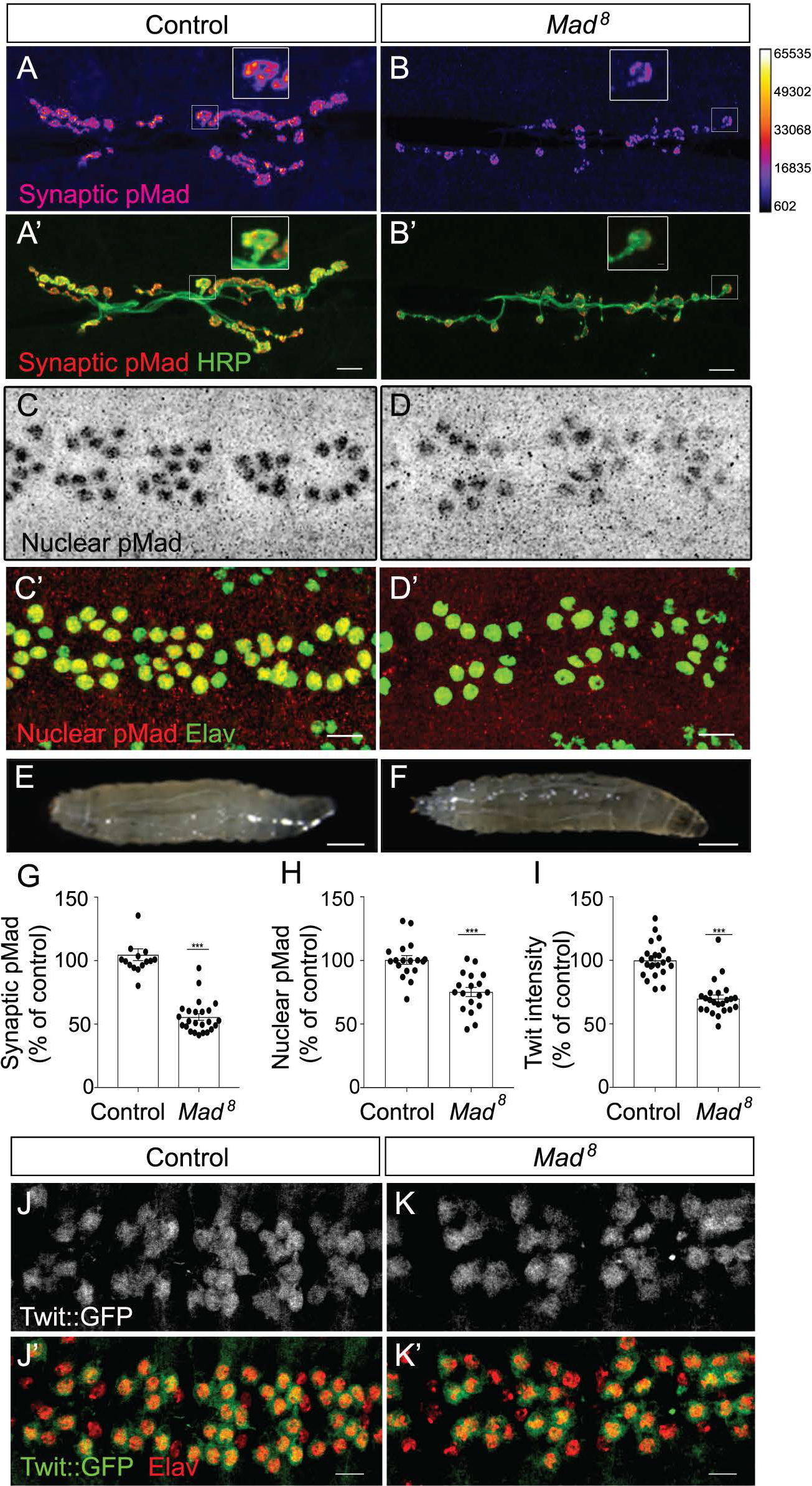
Disproportionate reduction of synaptic pMad in *Mad*^*8*^ mutants. (A-D) Confocal images of NMJ 6/7 boutons and ventral ganglia in control and *Mad*^*8*^ third instar larvae labeled for pMad (red) and HRP (green) (A-B) or Elav (green) (C-D). (E-F) Compared with control (E), fat body levels appear modestly reduced in *Mad*^*8*^ mutant larva (F). (G-I) Quantitative analysis of synaptic pMad (G), nuclear pMad (H) and Twit∷GFP (I). (J-K) Confocal images of ventral nerve cord of indicated genotypes showing motor neuron nuclei labeled with Twit∷GFP (green) and Elav (red). Twit∷GFP, a *MiMIC*-generated chimera, provides an additional read-out for the canonical BMP signaling. Genotypes: control (*Mi(MIC)twit*^*MI06552*^/+), *Mad*^*8*^(*Mad*^*8*^/*Df(2L)C28*, *Mi(MIC)twit*^*MI06552*^). Scale bars: 10 μm (A-D) and (J-K), 1 μm (details), 100 μm (E-F). Error bars indicate SEM. ***, p <0.0001

Since synaptic pMad mirrors the postsynaptic type-A glutamate receptors, we next examined the GluRIIA levels relative to total postsynaptic glutamate receptors, stained for GluRIIC, a subunit common for both type-A and type-B glutamate receptors (Marrus et al., 2004). Any disruptions in canonical BMP signaling reduce NMJ growth and produce NMJs with fewer boutons and a shrunken NMJ area (Marques and Zhang, 2006). Indeed, the 25% reduction in nuclear pMad in *Mad*^*8*^ mutants was accompanied by diminished HRP-labeled NMJ terminals and significantly decreased bouton area (to 57 ± 3% of control) (Figure 5A-C). Nonetheless, when normalized to HRP signals, synaptic GluRIIC levels appeared normal at *Mad*^*8*^ NMJs comparing to control (98 ± 6% of control, n= 25) (Figure 5A-B, quantified in D). In contrast, GluRIIA levels were significantly reduced at *Mad*^*8*^ NMJs (to 67 ± 5% of controls, n= 25).

**Figure 5.**
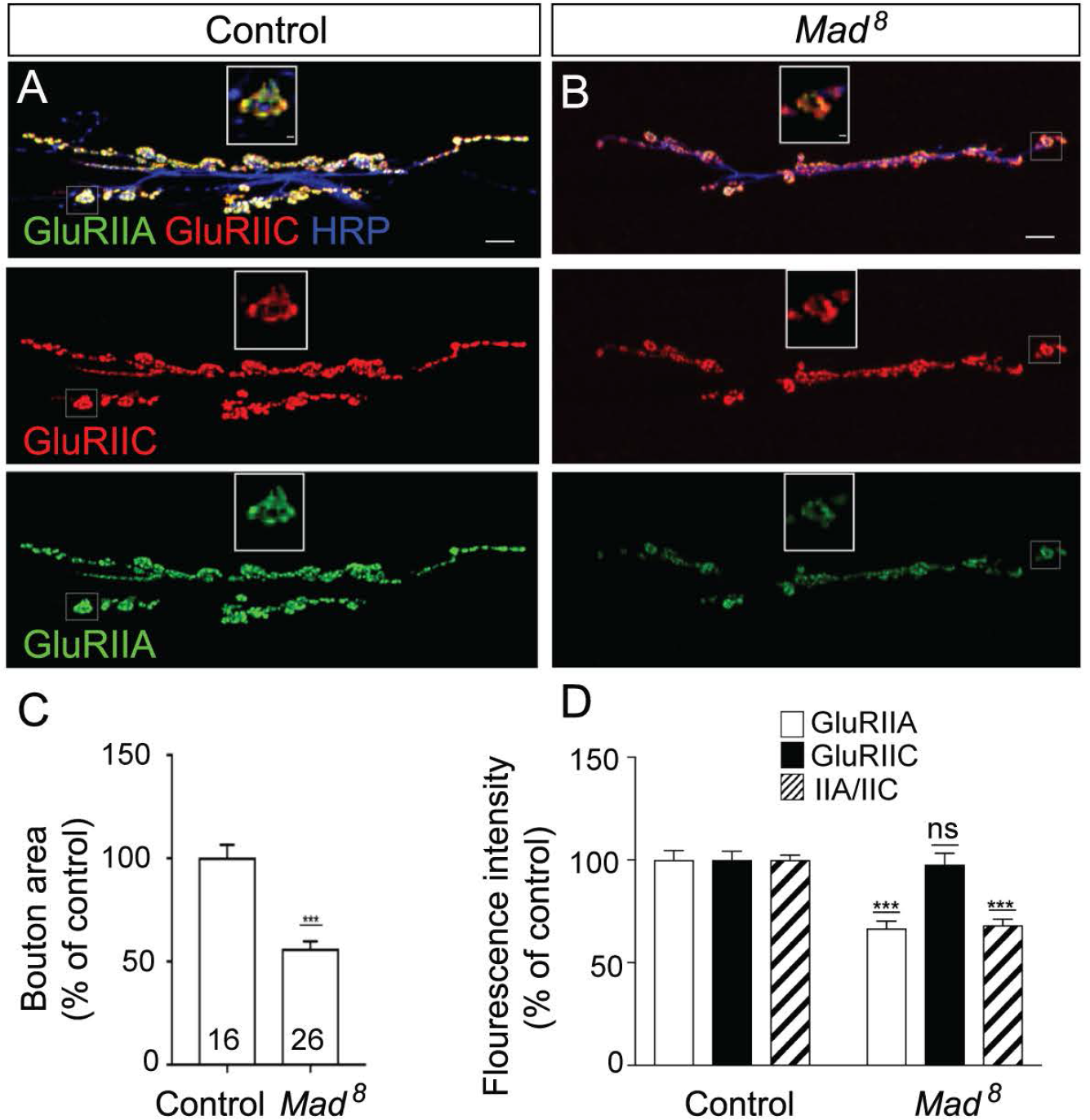
*Mad*^*8*^ mutants have altered postsynaptic iGluRs composition. (A-B) Confocal images of NMJ6/7 boutons from third instar larvae of control and *Mad*^*8*^ mutants labeled for GluRIIA (green), GluRIIC (red), and HRP (blue). *Mad*^*8*^ NMJs have reduced bouton area (C), diminished GluRIIA synaptic signals but normal GluRIIC levels (quantified in D). Scale bars: 10 μm, 1 μm (details). Error bars indicate SEM. ***, p <0.0001; ns, p >0.05.

Type-A receptors are determinants of the quantal size (the postsynaptic response to the release of single presynaptic vesicles) (DiAntonio et al., 1999). The reduced GluRIIA/GluRIIC ratio at *Mad*^*8*^ NMJs predicts that this mutant will have diminished quantal size. We tested this prediction by recording spontaneous junction currents from muscle 6 of control and *Mad*^*8*^ mutant third instar larvae. The mEJC amplitude was significantly reduced at *Mad*^*8*^ mutant NMJ (0.59 ± 0.03 nA in *Mad*^*8*^, n=11, vs 0.84 ± 0.04 nA in control, n=11, *p* =0.0003) (Figure 6A-C). Furthermore, the decay time constant was decreased in the *Mad*^*8*^ mutant (5.7 ± 0.23 ms for *Mad*^*8*^ vs 8.51 ± 0.40 ms for control, p < 0.0001), indicating a switch to faster desensitizing receptors at *Mad*^*8*^ NMJs (Figure 6D). Since type-B receptors desensitize much faster that type-A (DiAntonio et al., 1999), these data are consistent with the reduced relative levels of type-A receptors observed above (Figure 5). The mEJC frequency was also significantly reduced in *Mad*^*8*^ mutants (0.38 ± 0.08 Hz in *Mad*^*8*^ vs 1.01 ± 0.14 Hz in control; *p* = 0.001). This suggests a compound phenotype for *Mad*^*8*^ that may include additional contributions from targets of canonical BMP signaling, such as *twit* (Kim and Marques, 2012).

**Figure 6.**
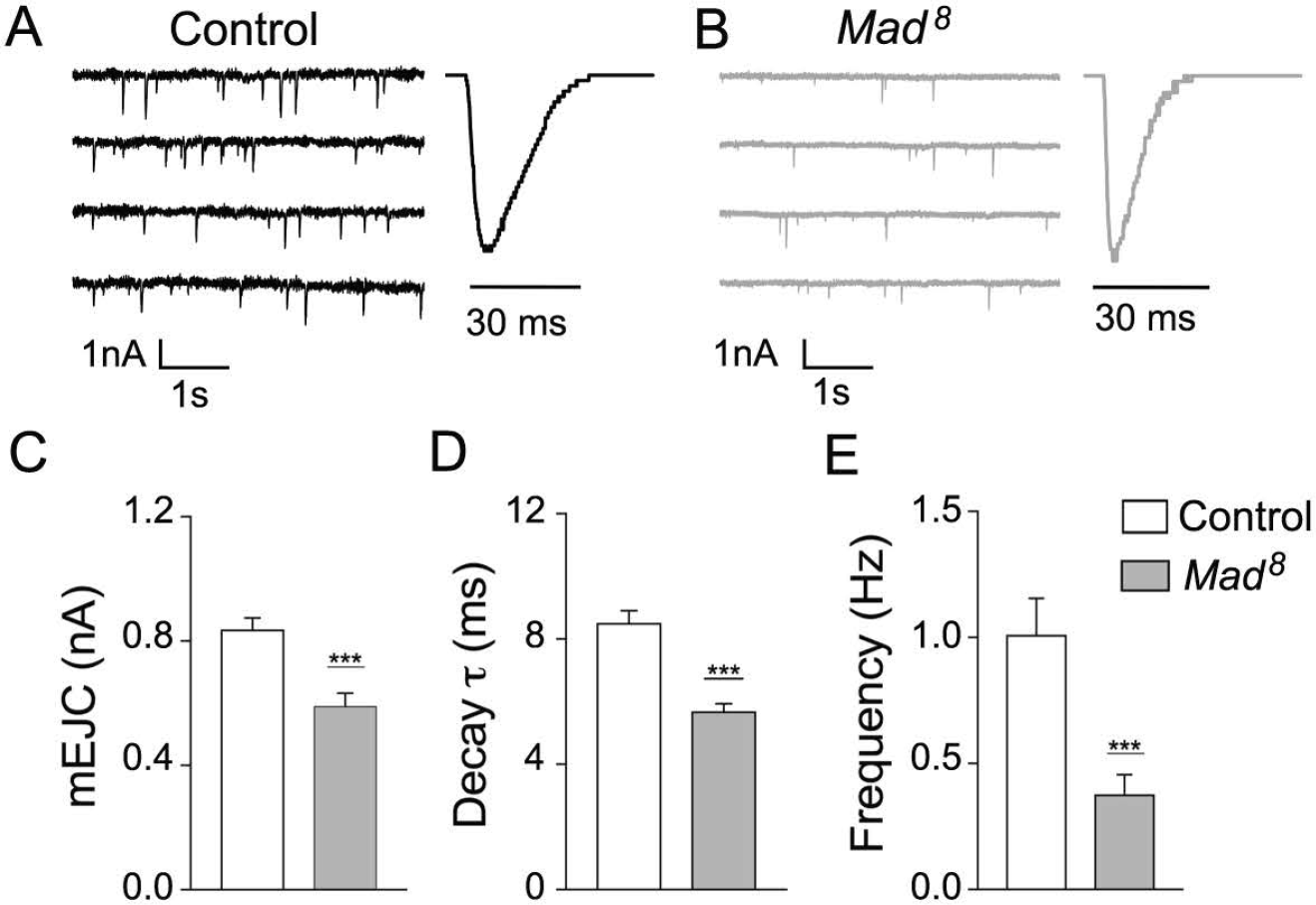
Electrophysiological deficits at *Mad^8^* NMJs. (A-B) Representative traces of spontaneous junction currents (left) and average mEJC traces (right) recorded at 0.5 mM Ca^2+^ from muscle 6, segment A3, of control (A) and *Mad*^*8*^ NMJs (B). Summary graphs showing the mean amplitude (C), decay time constant (D) and mean frequency (E) of mEJCs. Error bars indicate SEM. ***p< 0.0001.

### *In vitro* assays to measure pMad levels and accumulation at cell surfaces

To examine how *Mad*^*8*^ can primarily influence the synaptic pMad accumulation, we first analyzed its phosphorylation levels using a tissue culture-based signaling assay (Serpe and O’Connor, 2006). We transfected S2 cells with either wild-type or S359L Mad constructs and presented them with 0.1 nM Dpp, the fly ortholog of BMP2/4, which activates the BMP pathway. Dpp triggered a robust increase in pMad levels compared with the non-treated control (Figure 7 A-B); however, the pMad accumulation in cells transfected with Flag-Mad-S359L, called here Mad8, was only half of the wild-type control. This result could not be explained by a disruption of the Mad levels, as indicated by the internal control, Flag. Instead, this suggests that the S359L substitution severely impacts the ability of Mad8 to bind to and/or be phosphorylated by the Type I BMP receptors in response to BMP signaling.

**Figure 7.**
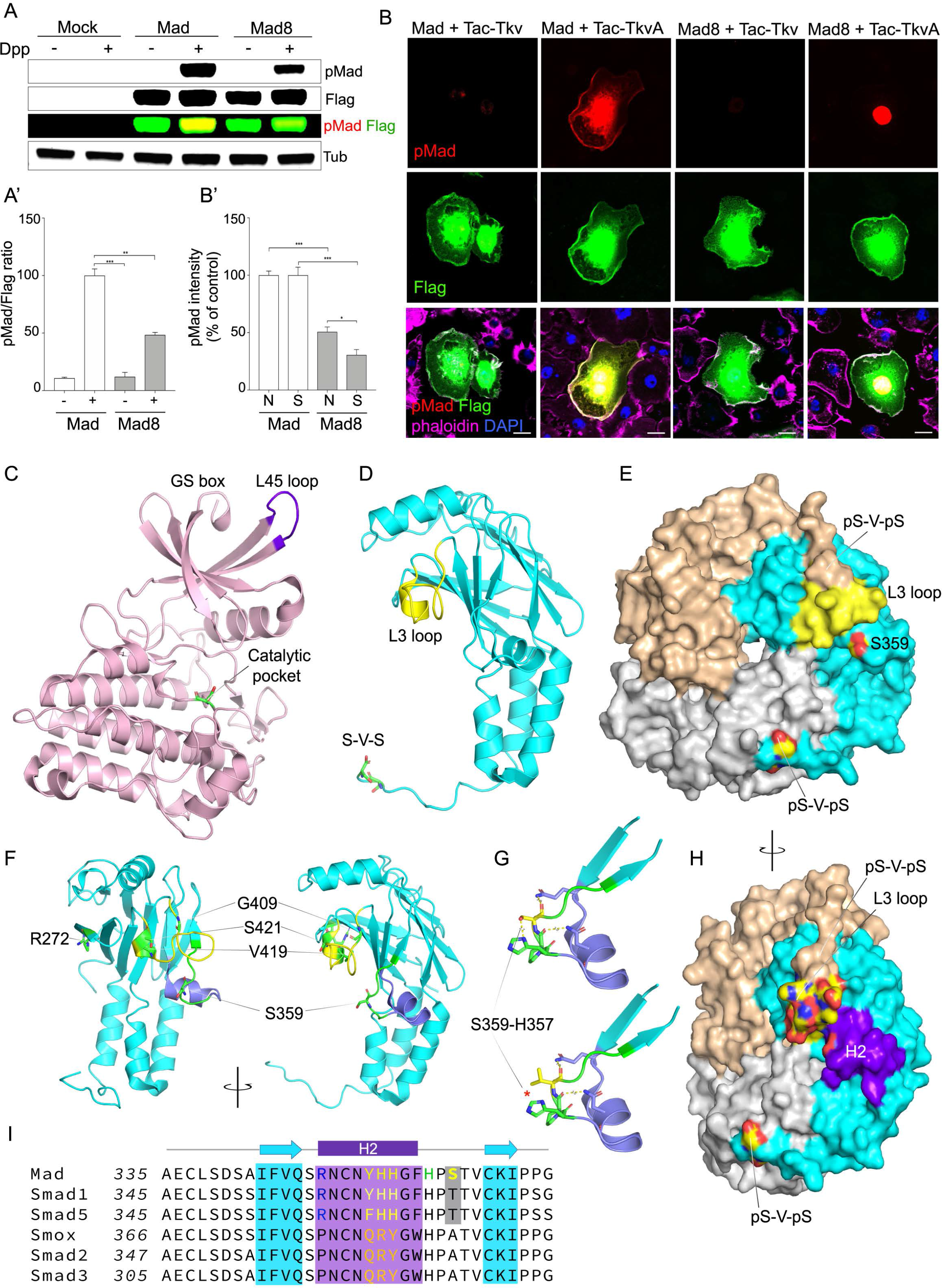
Biochemical analysis of Mad-Tkv interaction. (A) Western blot analysis of whole extracts from S2 cells expressing Flag-Mad and Flag-Mad-8 and treated with Dpp as indicated. Compared to control Mad, Mad-8 variant has significantly reduced pMad levels upon Dpp exposure. The pMad signals were normalized as the relative ratio pMad/Flag (A’). (B) Confocal images of S2 cells transfected with Flag-Mad variants and Tac-Tkv chimeras as indicated, spread on anti-Tac coated surfaces and labeled for pMad (red), Flag (green), actin (phalloidin-magenta), and DNA (DAPI-blue). The activated Tac-TkvA chimera induces nuclear pMad accumulation when co-transfected with Flag-Mad and, to a lesser extent, with Flag-Mad-8, as quantified in (B’). The pMad signals localize to cell surfaces only in Tac-TkvA/ Flag-Mad co-transfected cells. (C) Structure of the Type I receptor (PDM code 3TZM). The L45 loop (magenta) interacts specifically with R-Smads. The N-lobe of the receptor, including the GS box and L45, forms a docking surface for MH2 and positions the S-V-S C-tail of Mad in the catalytic pocket. (D-E) Structure of the MH2 domain of *Drosophila* Mad (PDM code 3DIT) shown as monomer (D) and trimer (E). The L3 loop (yellow) is engaged in exclusive interactions with either the L45 loop of the receptor or with the phosphorylated C-tail of another MH2 domain. (F) Map of MH2 Mad residues mutated in various *Mad* alleles. The two views of the structure are related by a 90° rotation around a vertical axis. (G) S359L in silica mutagenesis. S359 and adjacent peptide backbone form hydrogen bonds with H357 and residues on the H2 helix (purple); the S359L substitution breaks the hydrogen bonds with H357 (red asterisk) and introduces a bulky moiety, shifting the H2. (H) Lateral view of the Mad MH2 trimer showing the close proximity of the H2 helix (purple) to the charged L3 surface (colored by atoms). (I) Alignment of R-Smad sequences indicating class specific residues in the H2 region, including the S359 (yellow). Scale bars: 10 μm (B). Error bars indicate SEM. ***, p <0.0001; **, p <0.001; *, p <0.01

We have previously showed that synaptic pMad accumulates at active zones in puncta juxtaposing the postsynaptic densities (Sulkowski et al., 2016). This distribution near the presynaptic membrane together with the short life-time of pMad prompted us to propose that synaptic pMad (a) is generated locally by active BMP signaling complexes, and (b) remains associated with its own kinase, the activated Type I BMP receptor, in complexes anchored at the active zone via trans-synaptic interactions. Such signaling complexes, comprising ligands bound to tetrameric (type I and II) receptors and pSmads, have been also observed at vertebrate cellular junctions (Eom et al., 2011). Since *Mad*^*8*^ predominantly disrupts synaptic pMad, we hypothesize that Mad8 is deficient in binding to and/or remaining associated with BMP signaling complexes at the cell membrane. To test this possibility, we generated a Tac-Tkv chimera, where the Tkv intracellular domain was fused with the N-terminal and transmembrane domain of human Tac, the low affinity IL-2 receptor alpha chain (Ren et al., 2003). We also generated a Tac-activated Tkv chimera, referred here as Tac-TkvA, by introducing the Q199D substitution at the end of the GS box of Tkv; this substitution induces activation of Ser/Thr kinases within TGF-β Type I receptors (Wieser et al., 1995).

In the presence of Tac-Tkv, both Mad and Mad8 were detectable throughout cells, including at cell membranes (Figure 7B); however, no pMad immunoreactivity was visible, indicating an inactive Ser/Thr kinase in the Tac-Tkv chimera. Co-transfection of Mad with an activated Tac-TkvA chimera induced massive pMad accumulation, following the distribution of Flag signals throughout the cell. In contrast, only nuclear pMad signals were detectable for Mad8. Quantification of pMad signals showed a drastic reduction of nuclear pMad for Mad8 (50% less than in control, Mad transfected cells), and an even more profound reduction of pMad at cell membranes (to ~30% of control levels). These reduced *in vitro* pMad signals could reflect a disruption of the Tkv-Mad8 enzyme-substrate (E-S) interaction either by favoring the dissociation of the [E-S] complex or by increasing its energy of activation. Since pMad signals are particularly diminished at cell membranes, the S359L substitution likely favors the dissociation of Tkv-Mad8 complexes. This interpretation is consistent with our *in vivo* findings that *Mad*^*8*^ mutants disproportionally affect pMad accumulation at the active zones.

At this time, structures for Smad-Type I receptor complexes are not available, but site-directed mutagenesis and studies on related structures indicate that the L3 loops of R-Smads bind to the L45 loops of the receptors (Durocher et al., 2000; Huse et al., 1999)(Figure 7C-D); class-specific residues on both L3 and L45 confer specificity during signaling, *i.e.* BMP vs. Activin pathway. Also, the GS domain and N-terminal kinase lobe of the receptor appear to form an MH2 docking interface that positions the Smad SSXS C-tail adjacent to the ATP-binding pocket of the receptor. An opposing concave interface has been observed in all Smad MH2 domains, including the *Drosophila* Mad MH2. Once phosphorylated, pSmads have a high propensity to trimerize that favors their dissociation from receptors (Kawabata et al., 1998). The L3 loop of R-Smads has been implicated in mutually exclusive interactions with the Type I receptors or with the phosphorylated pS-X-pS tail of other Smads in trimeric complexes (Wu et al., 2001) (Figure 7E). This model is consistent with our experimental observations: Strong L3 mutations could effectively impair BMP signaling by disrupting both complexes, Mad-Tkv and Mad hetero-trimers, while moderate alleles may cause more limited disruptions of Mad-Tkv interaction.

S359, the residue mutated in Mad8, maps within the same concave cavity of MH2 and follows a highly conserved helix, H2 (Figure 7F-I). Inspection of this region indicates that S359 stabilizes the highly conserved H357 via hydrogen bonds. Additional hydrogen bonds connect negative charges of the peptide backbone with N349 and Q346. Together these interactions appear to anchor and stabilize the H2 helix. Similar to L3, the H2 helix contains class-specific residues: Y^352^HH in Smads of the BMP pathway, and QRY in the equivalent position in Smads of the Activin pathway (Figure 7I). The class-specific features of the H2 helix and its spatial orientation and proximity to the L3 loop suggest that H2 is a critical molecular determinant for the Smad-Type I receptor interaction. In this scenario, the S359L substitution in Mad8 disrupts the Mad-Tkv interaction by (1) replacing a polar interface residue with a bulky hydrophobic one, (2) destroying the hydrogen bond S359-H357, and (3) likely shifting the position of H2, and the YHH class-specific residues, relative to the Mad-Tkv interface. This is consistent with our *in vivo* findings that the S359L substitution induces a predominant reduction of synaptic pMad.

## DISCUSSION

Our comprehensive analysis of numerous D*rosophila Mad* alleles revealed a rich layer of complexity for this allelic series during different BMP signaling modalities and uncovered new molecular determinants that confer class-specificity during TGF-β signaling.

Using the *Drosophila* NMJ system, we have quantified for the first time the effect of *Mad* alleles on both canonical and non-canonical, Smad-dependent, BMP signaling pathways. *Drosophila* requires the canonical BMP pathway for NMJ growth. This pathway is triggered by muscle-derived BMPs which bind to their receptors on motor neurons and form active BMP signaling complexes that are endocytosed and retrogradely trafficked to motor neuron soma where they phosphorylate Mad (Marques and Zhang, 2006). The pMad accumulation in motor neuron nuclei quantitatively captures this canonical BMP response (Kim and Marques, 2010). In addition, pMad accumulates at synaptic terminals as a result of a non-canonical, but Smad-dependent BMP signaling modality, which sculpts postsynaptic composition as a function of activity (Sulkowski et al., 2014; Sulkowski et al., 2016). We have previously proposed that synaptic pMad is generated locally by active BMP signaling complexes confined to the active zone, a region protected from endocytosis (Sulkowski et al., 2016). This suggests that motor neurons could monitor synapse status then deploy the tightly controlled BMP receptors to (a) active zone, where they strengthen the synapse, or (b) outside the active zone, where they engage in canonical signaling to grow the NMJ.

As expected, strong *Mad* alleles drastically reduced both nuclear and synaptic pMad levels (Figure 1 and Table 2). In contrast, moderate *Mad* alleles reveal differential requirements for different signaling modalities. First, uniform reduction of Mad levels preferentially reduced local pMad, indicating that pMad accumulation at synaptic junction is highly sensitive to non-canonical signaling (Figure 2 and Table 2). This reduction may reflect (i) net levels of Mad within the motor neurons, presumably more abundant in the soma than in neurites, or (ii) the nature of the complexes, with Mad-BMPRI associating transiently in the soma but bound more stably at the synapses. Second, the prominent reduction of nuclear pMad in *Mad*^*1*^ mutants reinforces the idea that that the MH1, the DNA binding domain of Mad affected in this mutant, is critical for the nuclear accumulation of pMad and proper transcriptional control (Figure 3 and Table 2). Third, mutations in the L3 loop of the MH2 domain produced a set of moderate alleles (*Mad*^*9*^ and *Mad*^*11*^) with predominant deficits in synaptic pMad, but also produced two of the strongest *Mad* alleles (*Mad*^*7*^ and *Mad*^*10*^) (Figures 1 and 2). Since the L3 loop of R-Smads engages in mutually exclusive interaction with the type I receptors or with the phosphorylated C-tail of other Smads, we reason that strong L3 mutations may disrupt both sets of L3-dependent interactions, while moderate allele may elicit more limited disruption of Mad-Tkv interaction. Finally, the disproportionate deficits of synaptic pMad in *Mad*^*8*^ revealed a role for the H2 helix in the modulation of Mad-Tkv interaction. This highly conserved helix resides adjacently to the L3 loop and includes additional class-specific residues. The H2 features and the relative position to the MH2 concave cavity prompted us to propose that the H2 region is a critical determinant for the Smad-Type I receptor interaction and class specificity during signaling. H2 maps outside the Smad trimer interface (Figure 7H), but it could further interfere with Smad-dependent transcription via Smad/co-factor interactions. Because of our *in vivo* findings, we favor a role for H2 in modulation of the Mad-Tkv interaction. The H2 contribution may be direct, by shaping the Mad-Tkv interface, or indirect, via recruiting other protein(s) that may stabilize Mad-Tkv complexes at cell junctions.

Indeed, other proteins appear to contribute to the stabilization of BMP signaling complexes at specialized cell junctions. Previous studies in the chick neural tube showed that BMP signaling controls apicobasal polarity partly by enabling the pSmad1/5/8-dependent association of BMP signaling complexes with the PAR3-PAR5-aPKC complex at the tight junctions (Eom et al., 2011). Reduced Smad phosphorylation destabilizes the PAR complex and disrupts the tight junctions. In flies, loss-of-function disruptions of Bazooka(Par-3)-Par-6-aPKC complexes produced NMJs with significantly reduced number of boutons and increased levels of postsynaptic GluRIIA receptors (Ruiz-Canada et al., 2004). These phenotypes have been attributed to severe disruptions of microtubule stability, which may obscure a role for these complexes in modulation of BMP signaling.

In addition, posttranslational modifications of Mad and Tkv may strengthen or weaken their interaction and selectively modify nuclear and synaptic pMad. For example, Mad phosphorylation at S25 by Nemo (a) disrupts Mad association with Tkv at synaptic terminals, and (b) favors nuclear export of pMad (Merino et al., 2009; Zeng et al., 2007). In both flies and mammals, posttranslational modifications such as ubiquitination limit the activity of receptors and trigger degradation or deactivation of both Smads and receptors, keeping these signaling components in check (Dupont et al., 2009; Zhu et al., 1999). Interestingly, disruption of a number of components of the endocytic machinery elevate synaptic pMad and diminish nuclear pMad presumably by shifting receptors allocation from canonical BMP signaling, which requires retrograde transport, to local, non-canonical signaling (Heo et al., 2017; O’Connor-Giles et al., 2008; Vanlandingham et al., 2013).

Mis-regulation of BMP signaling is associated with many developmental abnormalities and disease states. Our study indicates that the H2 helix region is critical to local BMP signaling and suggests that mutations in this region may disrupt specialized tight junctions throughout the animal kingdom. We searched for human genetic variants in the H2 region using the MARRVEL database (model organism aggregated resources for rare variant exploration) (Wang et al., 2017). Interestingly, a Smad1 point mutation that changes a class-specific residue within H2, Y362C, was reported in the relative of a patient with colonic atresia, a congenital intestinal malformation that results in failure to pass meconium in newborns. Also, N361D substitution in Smad5 was found in a patient with malformation of the heart and great vessels. Finally, a N361-G365 deletion in Smad5 was reported in a patient with epileptic encephalopathy. While further studies are required to elucidate the functional impact of these genetic alterations on local BMP signaling, these variants underscore the relevance of the H2 helix for normal development and function.

## Acknowledgments

T.H.N, T.H.H, and M.S. were supported by Intramural Program of the National Institutes of Health, *Eunice Kennedy Shriver* National Institute of Child Health and Human Development, grants ZIA HD008914 and ZIA HD008869 awarded to M.S. S.N. was supported by NIH (OD024794). We are grateful to C.H Heldin for pMad antibodies. We thank Tom Brody and members of Serpe lab for comments and discussions on this manuscript. We also thank the Bloomington Stock Center at Indiana University for fly stocks and the Developmental Studies Hybridoma Bank at the University of Iowa for antibodies.

